# NF-κB-Dependent Transcriptional Regulation of Piezo1 Mediates Bacterial Clearance on Stiffened Lung Matrix

**DOI:** 10.1101/2025.11.06.687026

**Authors:** Erica M. Orsini, Adam M. Boulton, Susamma Abraham, Alyson Brown, Haley Ricci, Akash Ahuja, Bishnu Pant, Caitlin M. Snyder, Amanda Reinhardt, Amy H. Attaway, Ryan Musich, Lisa M. Grove, Mitchell A. Olman, Vidula Vachharajani, Rachel G. Scheraga

## Abstract

Respiratory pathogens, such as *Pseudomonas aeruginosa* damage the alveolar-capillary barrier leading to lung injury and stiffness. Lung stiffness is a key macrophage signal for bacterial clearance, but it remains unknown how stiffness-sensing mechanosensitive ion channels in macrophages are regulated during pneumonia. Macrophage Piezo1 is critical to bacterial clearance in experimental pneumonia *in vivo*; however, identification of putative matrix-derived signals and the mechanism of their effects remain to be determined. To our knowledge, our work is the first to show that during pneumonia, transcription of the mechanosensitive ion channel Piezo1 is increased in macrophages by the NF-κB transcription factor, p65, through its signaling adaptor protein, MyD88, leading to increased Piezo1 Ca^2+^ channel activity. Piezo1 mRNA abundance is increased in association with open chromatin at the Piezo1 promoter in macrophages. The enhanced level of Piezo1 increases the abundance of transcription factor EB (Tfeb) resulting in lysosome biogenesis and stiffness-dependent phagolysosome maturation, a critical step for macrophage bacterial clearance. Our data support the mechanism whereby transcription of macrophage Piezo1 is enhanced by p65 to augment bacterial clearance on an injured, stiffened lung matrix during pneumonia. Therefore, Piezo1 is a future therapeutic target against pneumonia-induced lung injury.

## INTRODUCTION

In the United States, pneumonia is a common cause of severe lung injury, known as acute respiratory distress syndrome (ARDS).^1,2^ *Pseudomonas aeruginosa* in an opportunistic bacteria which frequently leads to pneumonia-induced ARDS in individuals with immune suppression (e.g., sepsis) and/or structural lung disease (e.g., cystic fibrosis).^3-7^ *P. aeruginosa* virulence factors (e.g., flagellin and lipopolysaccharide [LPS]) and an overactive host response cause alveolar epithelial injury, alveolar collapse, fibrin deposition, and lung tissue stiffness.^8^ Increased lung stiffness (decreased lung compliance) is associated with a high mortality from ARDS (> 40%).^9-13^ As such, we seek to determine the molecular mechanisms in macrophages that mediate *P. aeruginosa* clearance in the context of an injured and stiffened lung matrix.

Lung macrophages are key effector cells which clear bacteria to limit pneumonia-induced lung injury and stiffness.^14,15^ Lung macrophage function is determined by microenvironmental cues, such as soluble bacterial (e.g., LPS, flagellin) and lung matrix (e.g., stiffness) signals.^16-19^ Soluble bacterial signals, known as pathogen-associated molecular patterns (PAMPs), bind to lung macrophage pattern recognition receptors (PRRs), such as the toll-like receptors (TLRs), to promote transcription of genes required for bacterial clearance functions (e.g., phagocytosis).^20,21^ *P. aeruginosa* virulence factors LPS and flagellin are PAMPs which bind to TLR4 and TLR5, respectively, leading to increased bacterial clearance.^22-26^ Activation of TLRs by PAMPs leads to recruitment of the signaling adapter protein MyD88 for activation of the pro-inflammatory transcription factor NF-κB. Activation of TLRs and various cytokine receptors (e.g., IL-1β and TNF-α) induces translocation of the NF-κB subunits (e.g., p65 [RelA], RelB, c-Rel, p52 and p50) to the nucleus for gene transcription.^27^ p65 is primarily responsible for pro-inflammatory transcriptional activity due to its C-terminal transactivation domain binding directly to DNA.^28^ p65 binds to gene regulatory elements (e.g., promoter and enhancer) for increased transcription of genes necessary for bacterial clearance. While our prior publications show that bacterial clearance is enhanced by a convergent signal from TLR4 activation and enhanced matrix stiffness, the signals that drive the essential physiological processes that underlie macrophage bacterial clearance remain incompletely defined.^17-19^

Our lab and others have investigated important macrophage functions that are initiated by mechanosensitive ion channels.^18,19,29^ Piezo1 is a ubiquitously expressed mechanosensitive cation channel that is activated by cell membrane tension leading to Ca^2+^ influx.^29,30^ Piezo1’s tension sensing feature is a consequence of its unique three-blade, propeller-like structure, making it distinct from other mechanosensitive ion channels, such as TRPV4.^31^ Piezo1 function can be regulated on multiple levels, including transcription, subcellular location, post-translational modifications (e.g., phosphorylation, glycosylation), and mechanical gating threshold.^32-36^ The regulation of macrophage Piezo1 and its functions important to bacterial clearance are not yet fully understood. We sought to determine how macrophages integrate bacteria-derived signals leading to enhanced Piezo1 Ca^2+^ signaling and augmented bacterial clearance in response to injured lung matrix. Our work shows for the first time that Piezo1 mRNA abundance is increased by NF-κB/p65 leading to increased lysosome biogenesis and phagolysosome maturation on stiffened lung matrix. This work furthers our understanding of macrophage-matrix interactions with clinical implications for a broad range of bacterial infections.

## METHODS

### Antibodies and Reagents

The following antibodies were used: Piezo1 (Proteintech, Cat. No. 15939-1-AP), Tfeb (Cell Signaling, Cat. No. 83010), IgG (Sigma, Cat. No. I5381-5MG), and AlexaFluor-488 (Invitrogen, Cat. No. 2382186). The following reagents were used: Yoda1 0-500 μM (Tocris, Cat. No. 5586), *P. aeruginosa* flagellin 250 ng/mL (InvivoGen, Cat. No. tlrl-pafla), LPS 100 ng/mL (Sigma Aldrich, Cat. No. L2630), BAY 11-7082 3 µM (Sigma-Aldrich Cat. No. 196870), LysoTracker 100nM (Invitrogen, Cat. No. L12492), and pHrodo Zymosan Particles 50 µg/mL (Invitrogen, Cat. No. P35364). Polyacrylamide gels (1, 8, and 25 kPa) were custom ordered from Matrigen Life Technologies (Bream, CA). The FLIPR5 Ca^2+^ assay kit (Cat. No. R8185) was purchased from Molecular Devices. Piezo1-targeted (Cat. No. SR423525) and control siRNA (Cat. No. SR30005) were purchased from Origene.

### Cell Culture, Nucleofection, and Immunoblot Analysis

All animal protocols were performed as approved by the Cleveland Clinic Institutional Animal Care and Use Committee (IACUC). Bone marrow-derived monocytes (BMDMs) were obtained from 8- to 12-week-old C57BL/6 wild-type (WT), MyD88^-/-^, Piezo1^fl/fl^, and Piezo1^LysMCre^ mice (IACUC #2624) and differentiated into macrophages using recombinant mouse M-CSF (50 ng/ml; PeproTech) as previously published.^19^ BMDMs were plated on fibronectin-coated (10 mg/mL) polyacrylamide gels of pathophysiologic range stiffnesses (1, 8, or 25 kPa) or tissue-culture plastic. BMDMs were given Piezo1-targeted (Origene, Cat. No. SR423525) or control siRNA (Origene, Cat. No. SR30005) and treated with nucleofection solution (Lonza, Cat. No. VCA-1003) prior to nucleofection with program D-032. Knock-down was confirmed by RT-qPCR and Ca^2+^ channel activity. Transcription factor EB (Tfeb) protein abundance was measured using immunoblot.

### Chromatin immunoprecipitation (ChIP), bulk RNA, global run on (GRO), and assay for transposase-accessible chromatin (ATAC) sequencing analysis and ChIP and real time quantitative polymerase chain reaction (RT-qPCR)

A search for sequenced chromatin bound to the transcription factor p65 in macrophages was conducted in the National Library of Medicine database.^37^ A p65 ChIP-seq dataset performed in BMDMs ± LPS for 30 minutes was selected for analysis (NCBI GEO No. GSE225833).^38^ p65 occupancy of promoter and enhancer regions for various macrophage mechanosensitive ion channels, including Piezo1, Piezo2, TRPA1, TRPC6, TRPV1, and TRPV4 was graphed. ChiP-qPCR samples were prepared as previously published.^39^ Briefly, chromatin from BMDMs stimulated ± LPS 100 ng/mL or *P. aeruginosa* flagellin 250 ng/mL for 1 hour was crosslinked using 37% paraformaldehyde. Cells were lysed and sonicated in a ChIP buffer with protease inhibitor (Thermo Scientific, Cat. No. 78440). An aliquot of whole cell lysis was reserved. p65 antibody (Cell Signaling, Cat. No. 6956) or IgG (Sigma, Cat. No. I5381-5MG) was added to the chromatin solution and rotated at 4 °C overnight. Protein A/G magnetic beads (Thermo Fisher, Cat. No. 88802) beads were added to the chromatin solution followed by column isolation (Zymo Research, Cat. No. D5205). RT-qPCR was performed on p65 or IgG pulldown chromatin solution and whole cell lysate using primers for the Piezo1 promoter (forward: AGTTACTCGCACTGTGCCC, reverse: GGGTCAGGTCTTTGGGAGTC) and enhancer (forward: TTCCTAGTAAGCAGCGGTGG, reverse: CTCTGGGTCGCTTTGTTCCA) regions. RT-qPCR was performed on our candidate mechanosensitive ion channel, Piezo1. BMDMs were stimulated ± LPS 100 ng/mL or *P. aeruginosa* flagellin 250 ng/mL for 6 hours. RNA was purified using column extraction (Qiagen, Cat. No. 74004). RNA was reverse transcribed into complementary DNA (Qiagen, Cat. No. 210210). RT-qPCR was performed using SYBR Green (Applied Biosystems, Cat. No. 4368706) on a QuantStudio3 (Applied Biosystems), as previously published.^18,19^ The following primers were used: Piezo1 (forward: TCTGTTCCTCACGCTGTTCC, reverse: CCAGCGCCATGGATAGTCAA) and Gapdh (forward: AGGTCGGTGTGAACGGATTTG, reverse: TGTAGACCATGTAGTTGAGGTCA). GRO-seq data were processed as described previously.^40^ For GRO-seq analysis, we parsed the genomic coordinates corresponding to the first 4,000 base pairs of the Piezo1 gene in BMDMs treated ± LPS. Analysis was performed on the summation of resulting peaks and graphed as normalized Piezo1 reads per million (NCBI GEO No. GSE140611).^40^ For bulk RNA sequencing, RNA was prepared from BMDMs using a column-based RNA isolation kit (Qiagen). RNA was sent to Novogene for sequencing and library preparation. Pathway analysis was performed using Ingenuity Pathway Analysis (Qiagen). To determine if Piezo1 mRNA abundance is increased during pneumonia *in vivo*, bulk RNA and ATAC sequencing data from mice with and without *P. aeruginosa* pneumonia was identified in the National Library of Medicine database (NCBI GEO No. GSE233206).^41^ Piezo1 normalized read counts from alveolar macrophages and lung tissue were quantified. Open chromatin along the Piezo1 promoter and enhancer was visualized using the University of California Santa Cruz Genome Browser (https://genome.ucsc.edu/).

### Piezo1 Abundance

Piezo1 cell surface abundance was measured in WT and MyD88^-/-^ BMDMs using flow cytometry (Attune NxT). BMDMs were stimulated ± *P. aeruginosa* flagellin 250 ng/mL for 6 hours. Cells were treated with Fc block (Biolegend, Cat. No. 101320) and stained with a Piezo1 primary antibody (Proteintech, Cat. No. 15939-1-AP) and AlexaFluor-488 secondary antibody (Invitrogen, Cat. No. 2382186). Next, a Sytox Blue (Invitrogen, Cat. No. S34857) live/dead stain was applied to the cells. Flow cytometry data was analyzed using FlowJo Version 10 (https://flowjo.com/). Gating strategy is outlined in **Supplemental Figure 1**. Piezo1 surface abundance was quantified as the frequency of AF-488 positive events. Total Piezo1 abundance was measured by immunofluorescence. WT and MyD88^-/-^ BMDMs were plated on an 8-well chamber slide (Ibidi) and stimulated ± *P. aeruginosa* flagellin 250 ng/mL for 6 hours. Cells were fixed using paraformaldehyde 4% for 20 minutes. Cells were treated with a Piezo1 primary antibody (Proteintech, Cat. No. 15939-1-AP) and AlexaFluor-488 secondary antibody (Invitrogen, Cat. No. 2382186). Slides were mounted using a DAPI containing media (ThermoFisher, Cat. No. P36935).

Cells were imaged using confocal microscopy (Leica) with three high-powered fields obtained per condition per biological replicate. Piezo1 abundance was quantified as integrated density/cell/high powered field using ImageJ (https://imagej.net/ij/).

### Piezo1 Ca^2+^ Channel Function

Piezo1 Ca^2+^ channel function was measured using the FLIPR 5 Ca^2+^ assay kit (Molecular Devices, Cat. No. R8185). BMDMs were stimulated by *P. aeruginosa* flagellin 250 ng/mL overnight. The Ca^2+^ loading dye was prepared and added to the cells according to the manufacturer instructions, as previously published.^19^ BMDMs were treated with varying concentrations of the Piezo1 specific agonist, Yoda1 (0-500 µM). Piezo1 Ca^2+^ channel activity in response to Yoda1 was quantified in relative fluorescence units (RFU) on the FlexStation 3 (Molecular Devices), as previously published.^18,19^

### Lysosome Biogenesis, Phagolysosome Maturation, and Bacterial Clearance

Lysosome biogenesis was measured using LysoTracker, as previously published.^42^ Briefly, BMDMs stimulated ± *P. aeruginosa* flagellin 250 ng/mL were treated with 100 nM LysoTracker for 30 minutes. Lysosome biogenesis (mean lysosome volume) was measured using confocal microscopy (Leica) and quantified using Volocity Visualization and Analysis Software (PerkinElmer). Phagolysosome maturation was measured using Zymosan A pHrodo bioparticles, as previously published.^17^ BMDMs were plated on polyacrylamide gels of pathophysiologic range lung stiffnesses (1 kPa = normal lung, 8-25 kPa = injured/stiffened lung) or standard tissue culture conditions (10^6^ kPa), and stimulated with 10^4-5^ colony forming units (CFU)/mL of a heat-inactivated clinical strain of *P. aeruginosa* (PAM 57-15) for 6 hours. Phagolysosome maturation was measured using the Flexstation 3 Microplate Reader (Molecular Devices) or confocal microscopy (Leica). BMDMs were given pHrodo bioparticles (for microplate reader 0.5 mg/mL and for confocal microscopy 50 µg/mL) for 45 minutes. Using the microplate reader, phagolysosome maturation was normalized using a plate blank for each stiffness (1, 8, 25, and 10^6^ kPa). As phagolysosome maturation cannot be a negative value, data points were baseline adjusted so the most negative value for each experiment was set at zero. For confocal microscopy, BMDMs were transferred to a chamber slide and fixed with 4% paraformaldehyde and mounted using DAPI containing media (ThermoFisher, Cat. No. P36935). Using confocal microscopy (Leica), phagolysosome maturation was measured using representative high-powered fields. Phagolysosome maturation was quantified as integrated density/cell/high powered field using ImageJ (https://imagej.net/ij/). To measure macrophage bacterial clearance, BMDMs were incubated with 10^4-5^ CFU of a live clinical strain of *P. aeruginosa* (PAM-57-15) for 30-60 minutes. BMDMs were washed with PBS and incubated with 100 ug/mL kanamycin for 2 hours. BMDMs were lysed with 0.1% Triton X and plated for CFU at 0 and 2 hours after kanamycin.

### Statistical Analysis

Statistical analysis was performed using GraphPad Prism 10 for Windows, GraphPad Software, San Diego, California USA (https://www.graphpad.com). Continuous variables, such as Piezo1 Ca^2+^ channel activity, Piezo1 abundance, and phagolysosome maturation, were tested for normality and outliers (Q-test with an α level of 0.01) and compared using a student’s t-test, Mann-Whitney U test, or analysis of variance (ANOVA) with tests for multiple comparisons (e.g., Dunnett’s and Šidák’s tests), as appropriate. The threshold for significance was set at p ≤ 0.05 (two-tailed). Data are presented as mean ± standard error of the mean (SEM).

## RESULTS

### Piezo1 Ca²⁺ channel activity is increased by matrix stiffness and further enhanced by *P. aeruginosa* flagellin through MyD88

To determine if the mechanosensitive ion channel Piezo1 is activated by pathophysiologic range lung stiffness, Piezo1 Ca^2+^ channel activity was measured upon plating WT BMDMs on varying stiffnesses in the pathophysiologic range or using standard tissue culture conditions (10^6^ kPa). The matrix stiffnesses recapitulated normal (1 kPa) and injured or stiffened (8-25 kPa) lung. Matrix stiffness increased Piezo1 Ca^2+^ channel activity in response to Piezo1-specific agonist, Yoda1, in a stiffness-dependent manner (**Figure 1A**). As *P. aeruginosa* is a common clinical respiratory pathogen which drives lung injury and stiffness, the effect of *P. aeruginosa* flagellin on Piezo1 Ca^2+^ activity was tested. *P. aeruginosa* flagellin further increased Yoda1-induced Piezo1 Ca^2+^ channel activity 10-fold, with a leftward shift of EC_50_ from ∼10 to ∼1 µM in WT BMDMs (**Figure 1B**). As MyD88 is an essential signaling protein downstream of flagellin activation of TLR5 in macrophages, Piezo1 Ca^2+^ channel activity was measured in WT and MyD88^-/-^ BMDMs ± flagellin.^43^ Yoda1-induced Piezo1 Ca^2+^ channel activity increased in WT (MyD88 sufficient) BMDMs in response to flagellin, an effect lost in MyD88^-/-^ BMDMs (**Figure 1C**). This suggests that signaling through MyD88 is required for increased Piezo1 Ca^2+^ channel activity. As Piezo1 channel activity may be related to enhanced Piezo1 protein abundance through MyD88, we stimulated WT and MyD88^-/-^ BMDM with *P. aeruginosa* flagellin and measured Piezo1 surface abundance by flow cytometry and total abundance by immunofluorescence (**Figure 1D-G**). Flagellin stimulation increased Piezo1 surface abundance by 25.5% (**Figure 1D-E**) and total abundance by 63% (**Figure 1F-G**) in WT BMDMs, an effect which was abrogated in MyD88^-/-^ BMDMs. These data show that Piezo1 Ca^2+^ channel activity is induced by matrix stiffness in a range that mimics injured lung (≥ 25 kPa) and is further increased by the *P. aeruginosa* virulence factor, flagellin, through MyD88.

**Figure 1.**
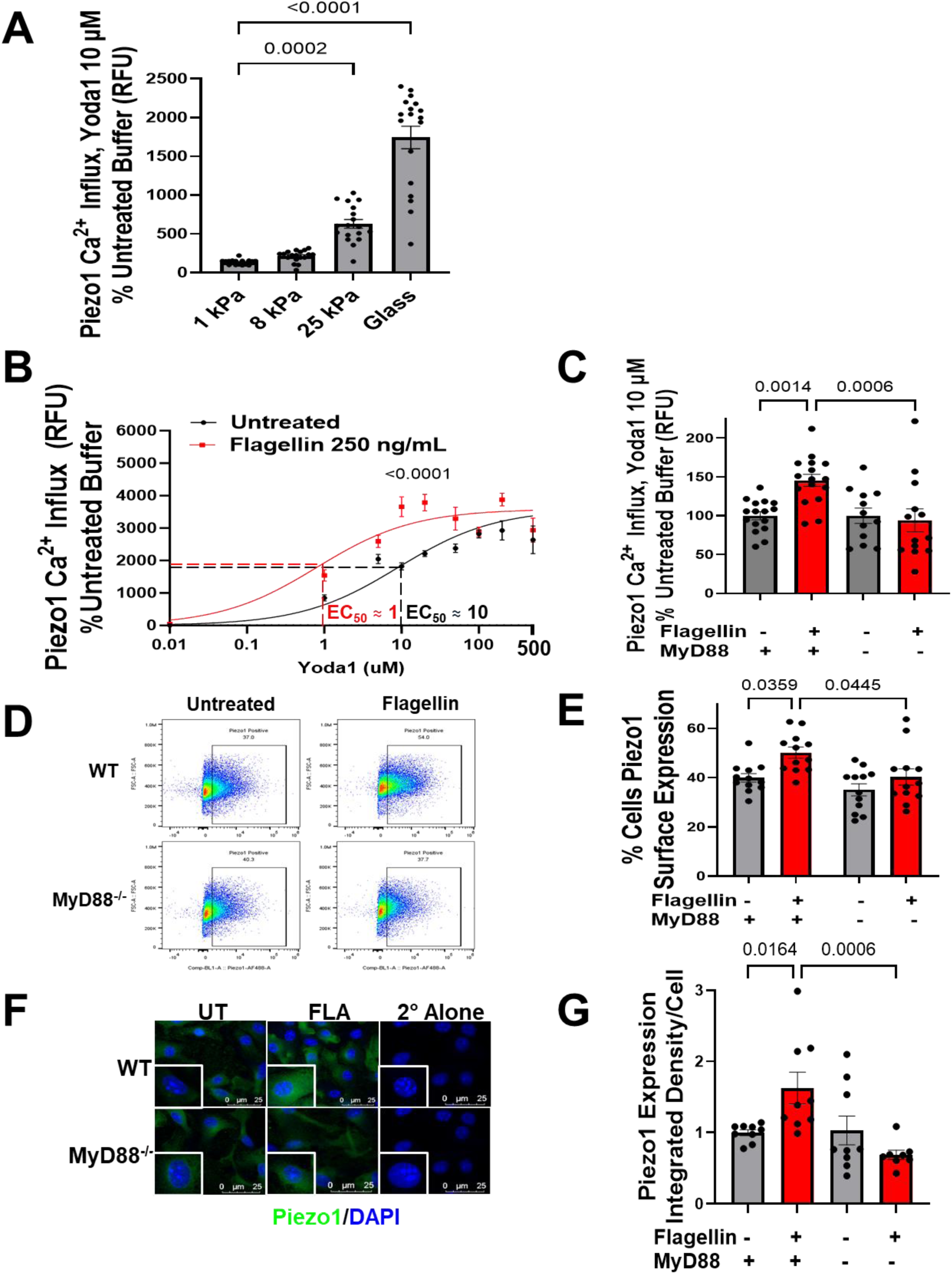
Piezo1 Ca²⁺ channel activity is increased by matrix stiffness and further enhanced by *P. aeruginosa* flagellin through MyD88. WT and MyD88^-/-^ BMDMs were plated on polyacrylamide gels of pathophysiologic range lung stiffness or standard tissue culture conditions and stimulated ± *P. aeruginosa* flagellin. (**A**) Piezo1 Ca^2+^ channel activity in response to Yoda1 requires increased matrix stiffness ≥ 25 kPa, similar to that seen in injured lung. (**B**) Flagellin increases Piezo1 Ca^2+^ activity above the Piezo1 agonist, Yoda1, alone, shifting the EC_50_ leftward from ∼10 to ∼1 μM on tissue culture plastic (10^6^). (**C**) The increased Piezo1 Ca^2+^ channel activity in response to flagellin was abrogated in MyD88^-/-^ BMDMs. (**D**) Piezo1 surface abundance by flow cytometry (quantified in **E**) and (**F**) total abundance by immunofluorescence (quantified in **G**), was increased upon flagellin stimulation in WT but not MyD88^-/-^ BMDMs. Data are presented as mean ± SEM. n = 3 - 6 biological replicates, comparisons by one-way ANOVA. p-values are as indicated.

### Piezo1 transcription is enhanced by NF-κB/p65 in response to LPS and flagellin

As flagellin enhances Piezo1 protein abundance through MyD88 and MyD88 is a regulator of NF-κB transcriptional activity, the role of NF-κB on Piezo1 transcription was interrogated using published chromatin immunoprecipitation sequencing (ChIP-seq) data (NCBI Geo No. GSE225833).^38,44^ As Piezo1 is not the only mechanosensitive ion channel in macrophages involved in lung injury and bacterial clearance, promoter and enhancer regions of Piezo1 and several additional macrophage mechanosensitive ion channels were investigated (**Figure 2**). As compared to the other mechanosensitive ion channels, Piezo1 had the highest predicted occupancy of p65 in its gene promoter and enhancer regions in LPS-treated macrophages (**Figure 2**). These data support that in LPS-treated macrophages, NF-κB/p65 is highly bound to the Piezo1 promoter and enhancer regions.

**Figure 2.**
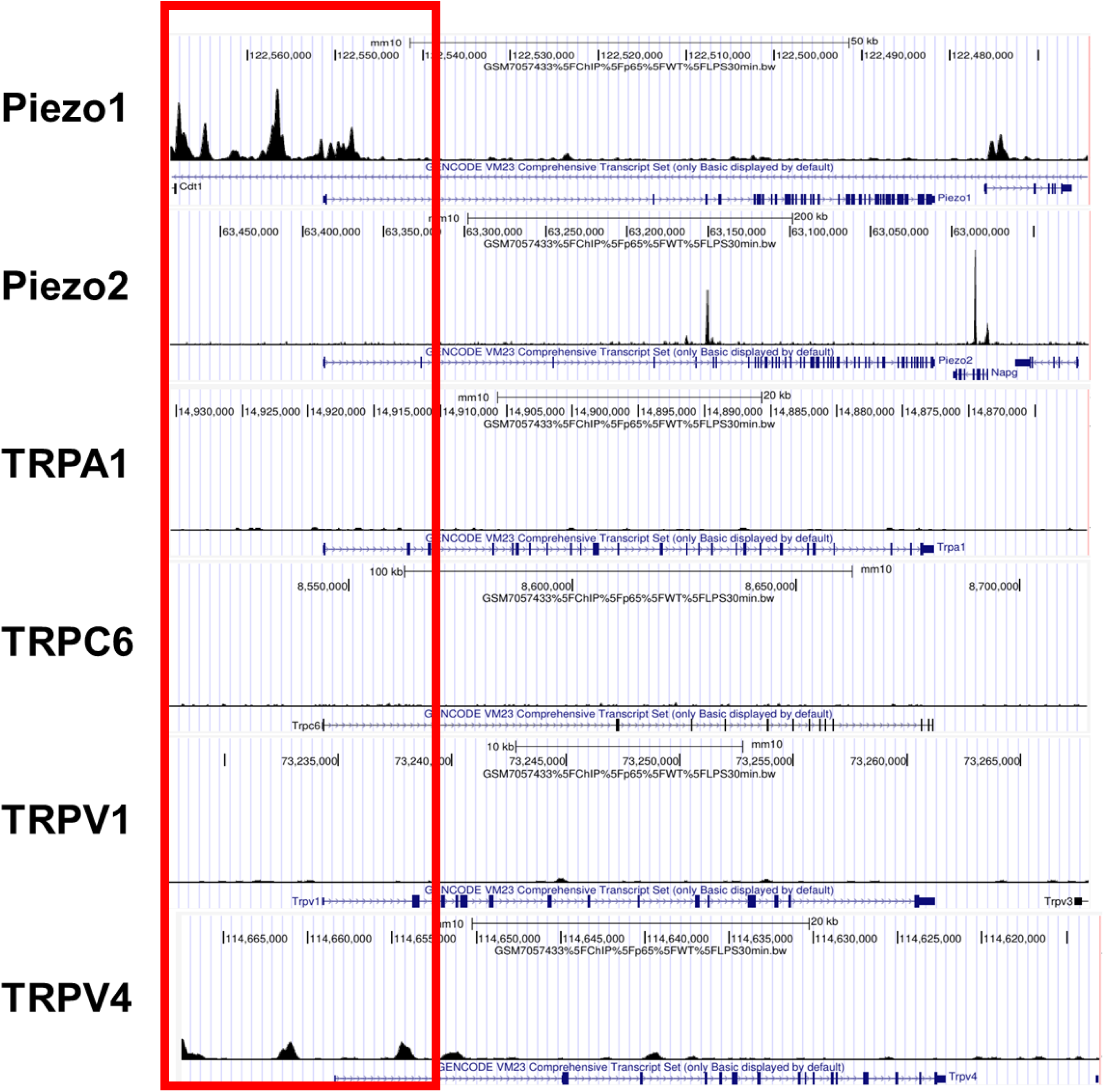
ChIP sequencing predicts Piezo1 mRNA abundance is regulated by the NF-κB transcription factor, p65. A published data set of chromatin bound to p65 from BMDMs stimulated ± LPS for 30 minutes was identified using NCBI GEO (GSE225833). We examined LPS-induced p65 occupancy at the promoter and enhancer regions of several macrophage mechanosensitive ion channels: Piezo1, Piezo2, TRPA1, TRPC6, TRPV1, and TRPV4 (red). The location of peaks represents genomic regions bound by NF-κB/p65 and the amplitude of peaks represents relative amount of NF-κB/p65 binding, as measured by reads per kilobase per million mapped reads, RPKM. Scale: 0-500 RPKM.

To validate the findings from the ChIP-sequencing dataset, p65 binding to the Piezo1 promoter (adjacent to the transcriptional start site) and enhancer (high p65 occupancy distal to the promoter) regions was measured in BMDMs upon *P. aeruginosa* virulence factors using ChIP-qPCR. p65 binding to the Piezo1 promoter was increased in response to LPS (by 2.3-fold, **Figure 3A**) and flagellin (by 1.5-fold, **Figure 3B**). NF-κB/p65 binding to the Piezo1 enhancer was similarly increased in response to LPS (by 2.8-fold, **Figure 3C**) and flagellin (by 2.4-fold, **Figure 3D**). The increased binding of p65 binding to the Piezo1 promoter and enhancer regions was associated with a ≥3-fold increase in Piezo1 mRNA in response to LPS (**Figure 3E**). The increased Piezo1 mRNA could be due to the production of nascent mRNA transcripts or enhanced mRNA stability.^45^ Analysis of a publicly available global run-on sequencing (GRO-seq) dataset (NCBI GEO No. GSE140611)^40^ revealed that the increase in Piezo1 mRNA abundance after LPS was due to an increase in nascent transcription (**Supplemental Figure 2**). In addition to LPS, another important *P. aeruginosa* virulence factor, flagellin, increased Piezo1 mRNA by ≥3-fold (**Figure 3F**). This effect was abrogated in MyD88^-/-^ BMDMs and in BMDMs treated with the NF-κB/p65 activity inhibitor, BAY 11-7082 (**Figure 3F**). These data demonstrate that p65 is an important regulator of Piezo1 transcription in macrophages in response to key *P. aeruginosa* virulence factors.

**Figure 3.**
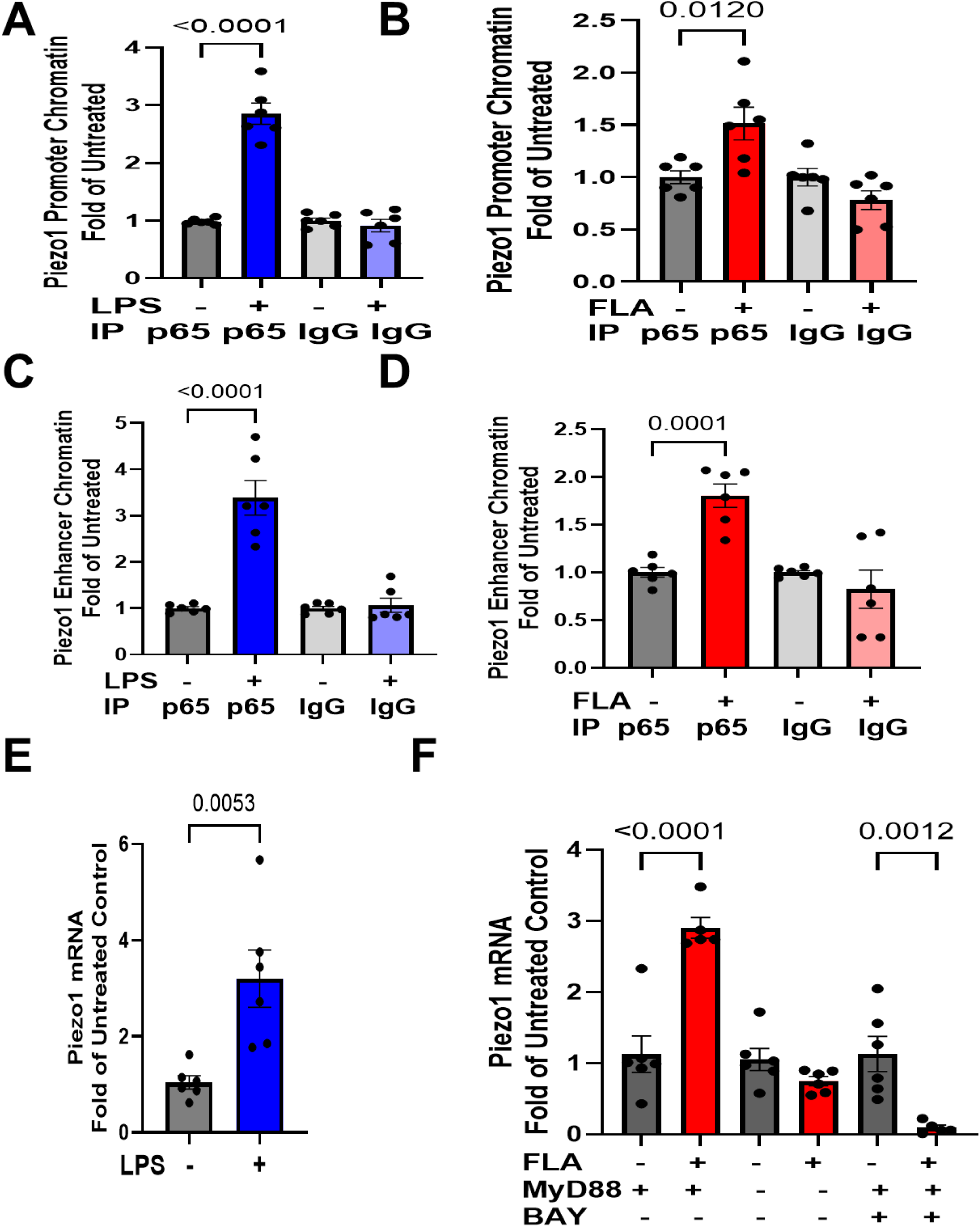
Piezo1 transcription is increased in response to LPS and flagellin through the NF-κB transcription factor, p65. WT and MyD88^-/-^ BMDMs stimulated ± LPS or flagellin (FLA) were plated on standard tissue culture conditions and ChIP- or RT-qPCR was performed. There was increased binding of p65 to the Piezo1 promoter in response to (**A**) LPS and (**B**) FLA and enhancer in response to (**C**) LPS and (**D**) FLA. Piezo1 mRNA transcription increased at 6 hours by > 3-fold in response to (**E**) LPS (100 ng/mL) and (**F**) FLA (250 ng/mL). Increased Piezo1 mRNA upon flagellin is abrogated in MyD88^-/-^ BMDMs and in WT BMDMs treated with the NF-κB inhibitor, BAY-11 7082 (3 μM, 6h). Data are presented as mean ± SEM. n = 3 biological replicates, comparison by t-test or one-way ANOVA, as appropriate. p-values are as indicated.

### Piezo1 regulates key macrophage bacterial clearance pathways and lysosome biogenesis

As NF-κB/p65 increases Piezo1 transcription upon exposure to *P. aeruginosa* virulence factors, the role of Piezo1 on flagellin-induced macrophage functions was queried. Bulk RNA sequencing (Novogene) was performed in BMDMs from Piezo1-sufficient (Piezo1^fl/fl^) and Piezo1-depleted (Piezo1^LysMCre^) mice upon *in vitro* exposure to *P. aeruginosa* flagellin. Piezo1 was found to regulate mRNA abundance of flagellin-induced phagocytosis pathways, including phagosome formation and maturation (**Figure 4A**). Detailed analysis reveals that Piezo1 mediates the flagellin-induced genes, including *Sqstm1*, *Aga*, and *Mcoln3*, as illustrated by a heatmap (padj < 0.05) (**Figure 4B**). *Sqstm1*, *Aga*, and *Mcoln3* are necessary for lysosome biogenesis, a critical step of macrophage bacterial clearance.^46^ These key lysosome genes are all downstream targets of the lysosome-regulating transcription factor, Tfeb.^47^ Thus, Tfeb protein abundance was measured in Piezo1^fl/fl^ and Piezo1^LysMCre^ BMDMs after flagellin by immunoblot. The flagellin-induced Tfeb abundance was reduced by 32% in the Piezo1-depleted BMDMs compared to the Piezo1-sufficient BMDMs after 1 hour which translated into a reduced lysosome biogenesis, as measured by Lysotracker (**Figure 4C-F**). Taken together, with the well described effects of Tfeb^47^, these data demonstrate that macrophage Piezo1 is required for bacterial clearance pathways in response to flagellin through enhanced Tfeb protein abundance and lysosome biogenesis.

**Figure 4.**
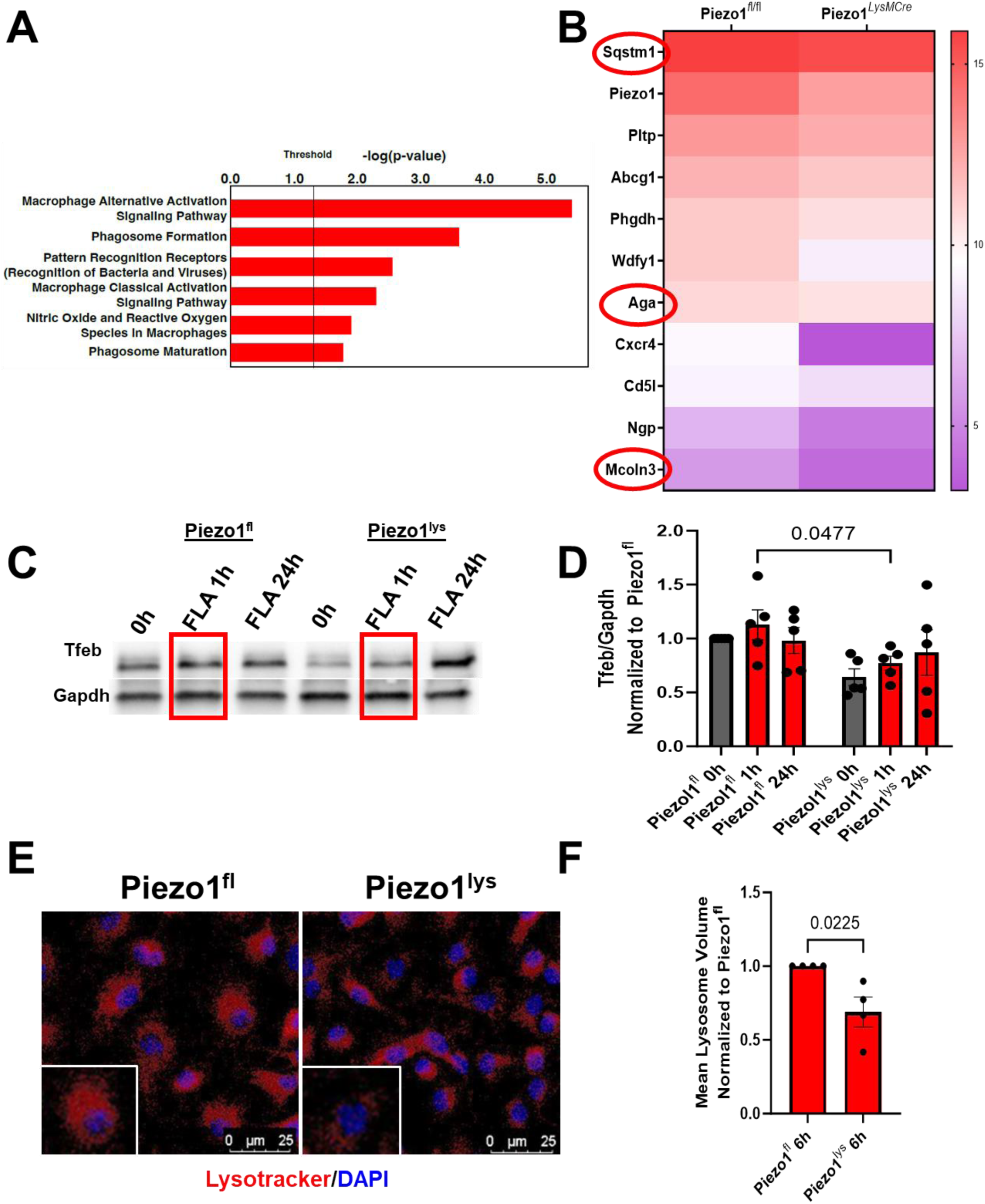
Piezo1 regulates key macrophage bacterial clearance pathways and lysosome biogenesis upon *P. aeruginosa* flagellin. Piezo1^fl/fl^ and Piezo1^LysMCre^ BMDMs were stimulated with *P. aeruginosa* flagellin for 6 hours using standard tissue culture conditions for bulk RNA sequencing, Tfeb protein abundance, and mean lysosome volume. (**A**) Significantly enriched bacterial clearance pathways (−log[p-value] ≥ 1.3) in RNAseq from Piezo1^fl/fl^ and Piezo1^LysMCre^ BMDMs stimulated with *P. aeruginosa* flagellin are shown. (**B**) Top bacterial clearance genes upregulated (padj < 0.05) in Piezo1^fl/fl^ compared to Piezo1^LysMCre^ BMDMs upon flagellin are shown (Log2-transformed normalized counts), including key targets of the Tfeb transcription factor, *Sqstm1*, *Aga*, and *Mcoln1* (red circle). (**C**) Piezo1^fl/fl^ BMDMs had increased Tfeb protein at one hour, as quantified in (**D**). (**E**) Piezo1 increases the mean lysosome volume in BMDMs upon *P. aeruginosa* flagellin (red), as quantified in (**F**). Data are presented as mean ± SEM. n = 3 - 5 biological replicates, comparisons by t-test or one-way ANOVA, as appropriate. Scale bars = 25 µm. Images were taken at 40x original magnification. p-values are as indicated.

### Phagolysosome maturation is stiffness-dependent through Piezo1 in *P. aeruginosa*-treated macrophages leading to enhanced bacterial clearance

As Piezo1 Ca^2+^ channel activity is augmented by matrix stiffness, the role of Piezo1 on macrophage phagolysosome maturation on injured lung stiffness was tested. Phagolysosome maturation was measured in WT BMDMs plated on polyacrylamide gels of pathophysiologic range lung stiffnesses (1 kPa = normal lung, 8-25 kPa = injured/stiffened lung), or standard tissue culture conditions (10^6^ kPa) upon exposure to heat-inactivated (80° C, 1 hour) *P. aeruginosa*. Phagolysosome maturation, as assessed by pHrodo, increased > 5-fold from 1 kPa (i.e., normal lung) to 25 kPa (i.e., injured lung), suggesting phagolysosome maturation is stiffness-dependent (**Figure 5A**). Next, to determine if phagolysosome maturation requires Piezo1, phagolysosome maturation was measured in *P. aeruginosa*-treated macrophages from Piezo1 sufficient and depleted mice on injured lung stiffness (25 kPa). First, the expected reduction of Piezo1 Ca^2+^ channel activity in Piezo1-depleted macrophages (Piezo1^LysMCre^) and after Piezo1-targeted siRNA treatment (> 50% knockdown mRNA) was confirmed (**Supplemental Figure 3**). Phagolysosome maturation was reduced by 29% in Piezo1-depeleted (Piezo1^LysMCre^) BMDMs compared to Piezo1-sufficient (Piezo1^fl/fl^) BMDMs and reduced in Piezo1-targeted siRNA treated BMDMs by 56% compared to control siRNA treated BMDMs (**Figure 5B-E**). Finally, bacterial clearance of live *P. aeruginosa* was reduced by 38.2% in Piezo1-depleted (Piezo1^LysMCre^) compared to Piezo1-sufficient (Piezo1^fl/fl^) BMDMs (**Figure 5F**). These data indicate that macrophages require matrix stiffness sensing through Piezo1 for phagolysosome maturation in response to *P. aeruginosa* virulence factors for effective bacterial clearance.

**Figure 5.**
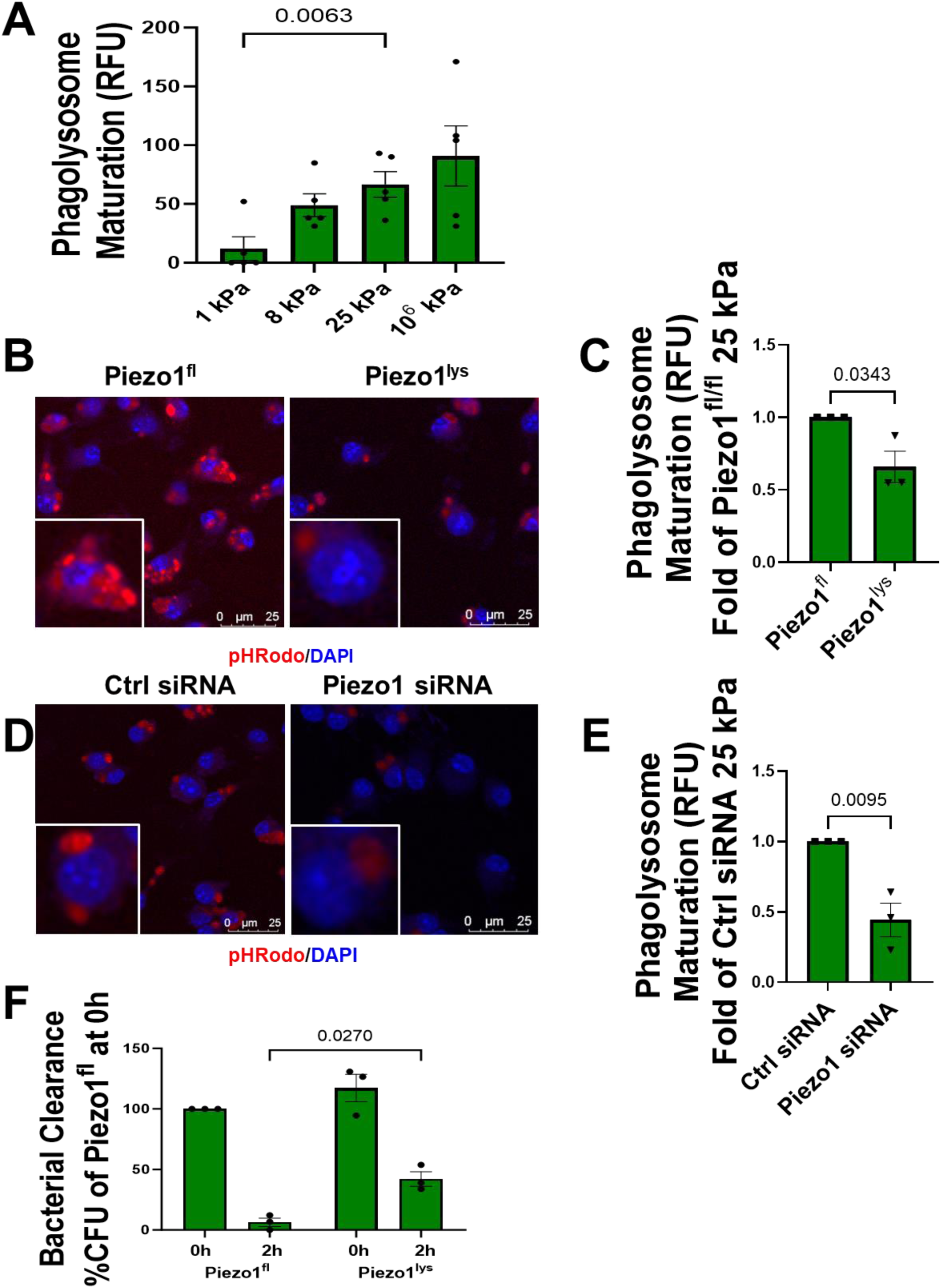
Phagolysosome maturation is stiffness-dependent through Piezo1 leading to enhanced bacterial clearance of *P. aeruginosa*. WT, Piezo1^fl/fl^, and Piezo1^LysMCre^ BMDMs were treated with heat-inactivated and live *P. aeruginosa* (green). Piezo1 Ca^2+^ channel activity, phagolysosome maturation, and bacterial clearance were measured on polyacrylamide gels of pathophysiologic range lung stiffness (1 kPa: normal lung, 8-25 kPa: injured lung) and standard tissue culture conditions (10^6^ kPa). (**A**) Phagolysosome maturation was measured using pH-sensitive fluorescent pHrodo bioparticles in *P. aeruginosa*-treated WT BMDMs on pathophysiologic-range matrix stiffness (1, 8, 25 kPa), n = 5. Phagolysosome maturation is stiffness-dependent and requires matrix stiffness which resembles injured lung (25 kPa). Phagolysosome maturation is reduced upon downregulation of Piezo1 by (**B**) Cre recombinase, as quantified in (**C**), or (**D**) siRNA, as quantified in (**E**), n = 3. (**F**) Bacterial clearance was measured using colony forming units (CFU) of *P. aeruginosa*. Bacterial clearance is enhanced in Piezo1-sufficient (Piezo1^fl/fl^) BMDMs, n= 3. Data are presented as mean ± SEM. Comparisons by one-way ANOVA or t-test, as appropriate. Scale bars = 25 µm. Images were taken at 40x original magnification. p-values are as indicated.

### Piezo1 is upregulated in alveolar macrophages after *P. aeruginosa*

As our *in vitro* data show that flagellin and lung stiffness conspire to upregulate Piezo1 activity, the relevance of our findings to *P. aeruginosa* pneumonia was investigated. Using a published RNA-seq and ATAC-seq dataset (GSE233206)^41^ from *P. aeruginosa*-infected mouse alveolar macrophage cell line and a C57BL/6 mouse model of *P. aeruginosa* pneumonia, transcriptional regulation of Piezo1 after *P. aeruginosa* was assessed. Piezo1 mRNA transcripts increased 64% in mouse alveolar macrophages after *P. aeruginosa*, compared to controls (**Figure 6A**). In contrast, whole lung Piezo1 transcriptional levels were not different ± *P. aeruginosa* pneumonia, suggesting that Piezo1 mRNA upregulation is unique to macrophages (**Figure 6B**). Next, to determine the mechanism by which Piezo1 mRNA abundance is increased in alveolar macrophages after *P. aeruginosa*, increased open chromatin accessibility along the Piezo1 promoter and enhancer regions of alveolar macrophages was assessed. There is increased open chromatin along the Piezo1 promoter and enhancer regions in alveolar macrophages after *P. aeruginosa* (**Figure 6C-D**). These data support that Piezo1 mRNA abundance is increased in alveolar macrophages following *P. aeruginosa*.

**Figure 6.**
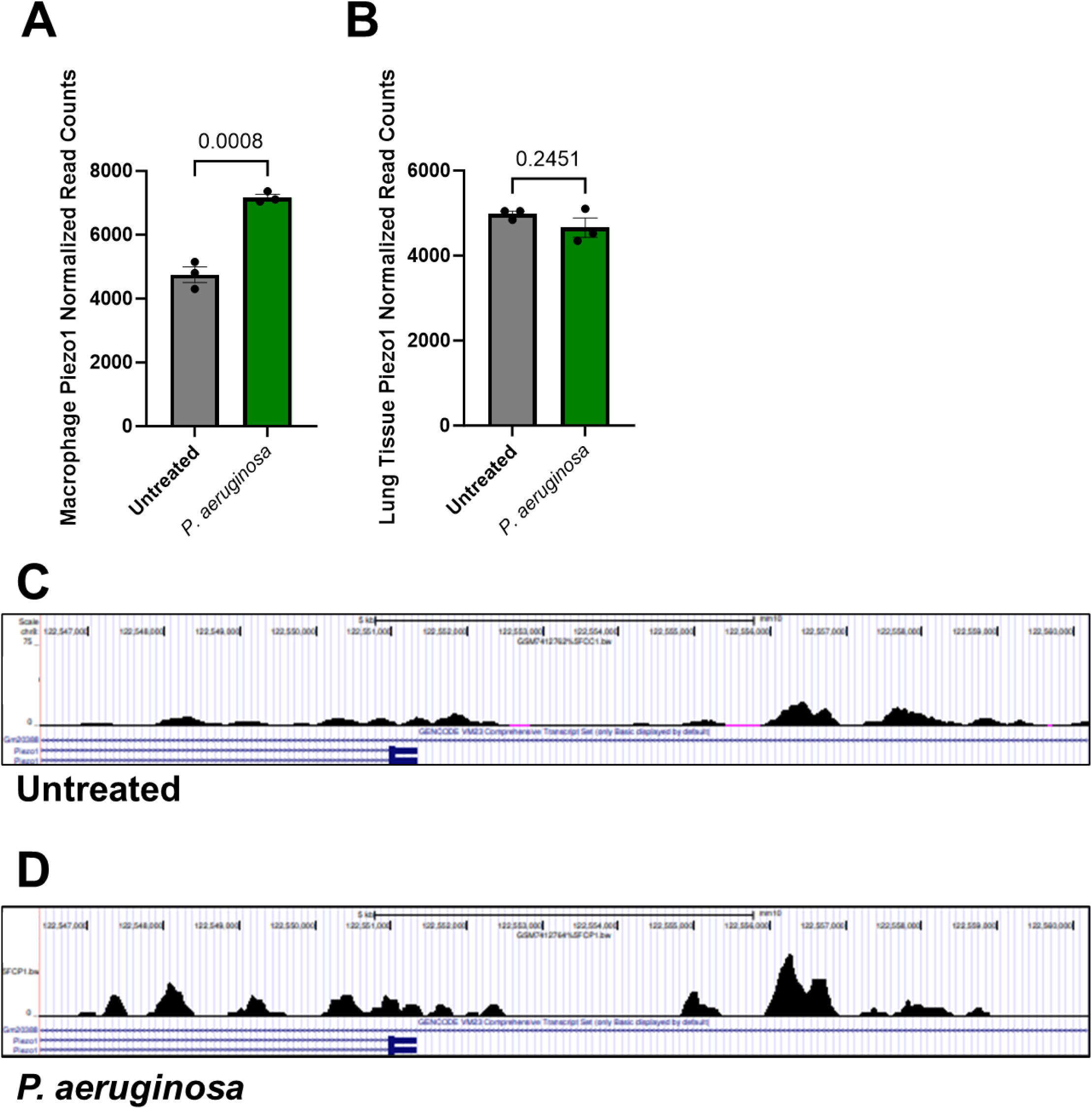
Piezo1 is upregulated in alveolar macrophages after *P. aeruginosa*. A published dataset containing RNA-seq and ATAC-seq from alveolar macrophages and lung tissue after *P. aeruginosa* was identified using NCBI GEO (GSE233206). Piezo1 normalized read counts from alveolar macrophages and lung tissue were quantified (n = 3 biological replicates/group). RNA-seq data reveal that Piezo1 is increased in (**A**) alveolar macrophages and not in (**B**) whole lung tissue after *P. aeruginosa*. Representative genome tracts of the Piezo1 promoter and enhancer reveal increased open chromatin in infected alveolar macrophages compared to controls (**C & D**). Scale: 0 - 75 reads per kilobase per million mapped reads, RPKM. Data are presented as mean ± SEM. p-values are as indicated.

## DISCUSSION

Building on existing literature, our current work shows a previously unrecognized link between the transcription factor NF-κB/p65 and transcriptional regulation of the mechanosensitive ion channel Piezo1. Specifically, we show that NF-κB/p65 binds to the Piezo1 promoter and enhancer regions leading to a MyD88-dependent increase in Piezo1 transcription, protein abundance, and Ca^2+^ channel activity in macrophages upon LPS and flagellin stimulation. Piezo1 regulates key bacterial clearance pathways and enhances protein abundance of lysosome-regulating transcription factor, Tfeb, thus increasing lysosome biogenesis. As a result, macrophage Piezo1 promotes phagolysosome maturation on an increased matrix stiffness which mimics injured lung (25 kPa) leading to bacterial clearance of *P. aeruginosa*. As highlighted in our prior work, lung stiffness is an important signal for macrophage bacterial clearance during pneumonia.^18,19,29^ However, until now, little was known about the mechanism whereby mechanosensitive ion channel transcription, abundance and function were regulated during pneumonia. As pneumonia induced-lung stiffness causes respiratory failure and death, it is critical that macrophages can effectively detect and respond to pneumonia-induced lung stiffness for rapid bacterial clearance and pneumonia resolution.^9,11,12^ Therefore, our findings provide critical insight into the relationship between pneumonia, transcriptional regulation of the macrophage mechanosensitive ion channel, Piezo1, and bacterial clearance.

Herein, we show that TLR activation by *P. aeruginosa* virulence factors (LPS and flagellin) augment Piezo1 abundance and Ca^2+^ channel activity which mediates enhanced phagolysosome maturation on stiffened lung matrix. However, increased Piezo1 abundance secondary to NF-κB/p65-dependent transcription is not the only mechanism to increase Piezo1 Ca^2+^ channel activity during infection. Others have also shown that LPS can augment Piezo1 Ca^2+^ activity through direct TLR4-Piezo1 binding.^48^ Furthermore, macrophage Piezo1 promotes stabilization of the transcription factor HIF1α to augment bacterial clearance during *P. aeruginosa* pneumonia.^29^ In addition to LPS, several other microenvironmental cues found in the infected lung drive increased Piezo1 mRNA abundance, including matrix stiffness, pro-inflammatory cytokines, and hypoxia.^32,49-51^ How these mechanical and inflammatory signals converge on Piezo1 to enhance macrophage function and thereby mediate resolution of pneumonia is an exciting area for future research.

Lung matrix stiffness (reduced compliance) contributes to respiratory failure and death from pneumonia-induced ARDS.^9,11,12^ Our lab and others have shown that mechanosensitive ion channels Piezo1 and TRPV4 are important for bacterial clearance during pneumonia *in vivo*, highlighting the importance of lung matrix stiffness-initiated signals to macrophage function.^18,19,29^ Our findings are supported by others’ work which also shows the importance of Piezo1 and TRPV4 to macrophage function during lung diseases, such as idiopathic pulmonary fibrosis (IPF) and ventilator-induced lung injury (VILI).^52-54^ Currently, the role of crosstalk between macrophage mechanosensitive ion channels Piezo1 and TRPV4 to coordinate bacterial clearance during pneumonia is poorly understood. In other cell types, crosstalk between Piezo1 and TRPV4 is an important contributory mechanism towards the pathobiology of several diseases, including pancreatitis, osteoarthritis, and in inflammation-induced vascular permeability.^55-58^ For example, in endothelial cells, activation of Piezo1 by shear stress leads to phospholipase A2 production of arachidonic acid (AA). AA actives TRPV4 leading to sustained Ca^2+^ influx which drives disruption of adherens junctions, actin remodeling, and vascular permeability, an important driver of lung injury.^57^ These findings highlight the need for cell-type specific research into macrophage Piezo1-TRPV4 crosstalk for stiffness-induced bacterial clearance.

We show that enhanced Piezo1 Ca^2+^ channel function in response to pneumonia-induced matrix stiffness controls phagolysosome maturation. However, it remains unknown how persistent exposure to *P. aeruginosa* virulence factors, such as during Cystic Fibrosis (CF) chronic lung infection, affects macrophage Piezo1 abundance and function.^59-62^ During early infection, p65/p50 heterodimers bind to gene promoters and enhancers. As a counterregulatory mechanism, p65 with its transcriptional activating domain (TAD) is degraded, leaving p50/p50 homodimers, which lack transcriptional activity, to bind to gene promoters, contributing in part to a suppressed response to LPS, known as endotoxin tolerance.^28,63^ Future work is needed to determine how chronic lung infections affect Piezo1 abundance and bacterial clearance in lung macrophages. Additionally, it is unclear how other ion channels, such as CF transmembrane regulator (CFTR) influence Piezo1 mRNA abundance during chronic lung infection. Therefore, Piezo1 abundance and function in chronic lung infection, such as CF chronic lung infection, is an important area for future research.

Our work has uncovered the novel finding that Piezo1 augments protein abundance of the lysosome-regulating transcription factor, Tfeb, lysosome biogenesis, and phagolysosome maturation in macrophages. As Tfeb self-regulates its abundance in a positive feedback loop, these data suggest that the mechanosensor, Piezo1, is required for Tfeb activity.^64^ These findings align with prior work on Piezo1’s control of lysosome function through Ca^2+^/calmodulin signaling and cytoskeletal regulation (e.g., Rho GTPase).^48,65^ Current evidence suggests that Piezo1 regulates lysosomal trafficking and exocytosis from the plasma membrane. However, whether Piezo1 is located on the lysosome membrane is an exciting area for future research. Lysosome Ca^2+^ channels are critical for lysosome trafficking, fusion, and acidification and several, including TRPML1, TRPML3, and TPC2 are mechanosensitive.^66-68^ Proteomics datasets have revealed low levels of Piezo1 on the lysosome membrane.^69^ However, whether Piezo1 regulates lysosome function on the lysosome membrane in response to intracellular mechanical cues is a fascinating area for future investigation.

Although our work significantly adds to the scientific field, we understand that our study has several limitations. We focused on transcriptional regulation of Piezo1 by NF-κB/p65, supported by our analysis of the p65 ChIP-seq omics data. However, it remains unknown if additional transcription factors promote Piezo1 abundance during *P. aeruginosa* pneumonia. In addition, this manuscript examines the role of Piezo1 on macrophage bacterial clearance on injured lung stiffness (25 kPa). However, our confirmatory ChIP experiments were performed on standard tissue culture conditions (10^6^ kPa), which are supraphysiologic. As matrix stiffness increases p65 transcriptional activity, future work will perform ChIP on pathophysiologic range of lung stiffnesses (1-25 kPa) to demonstrate the stiffness-dependency of Piezo1 transcriptional regulation by NF-κB/p65.^70,71^ Although NF-κB signaling is important to the development of pneumonia-induced lung injury, our work does not examine the effect of Piezo1 activation on the NF-κB pathway.^72^ However, emerging evidence suggests that activation of Piezo1 increases NF-κB transcriptional activity.^30^ Further understanding of the relationship between Piezo1 and NF-κB activation will provide insight into the effect of Piezo1 on lung injury during pneumonia. Our studies focused on increased Piezo1 abundance by the TLR5 agonist *P. aeruginosa* flagellin. However, flagellin’s activity is not entirely limited to TLR5. In addition to TLR5, flagellin activates the NLRC4 inflammasome.^73^ Therefore, it remains unknown how inflammasome activation affects Piezo1 abundance and function, although the literature supports that Piezo1 can promote inflammasome activation.^74^

In conclusion, the macrophage mechanosensitive ion channel Piezo1 is transcriptionally regulated by NF-κB/p65 in response to *P. aeruginosa* leading to enhanced Tfeb abundance, lysosome biogenesis, and bacterial clearance on injured, stiffened lung matrix (**Figure 7**). Lung stiffness is a critical signal to macrophages for bacterial clearance during pneumonia. Therefore, Piezo1 is a potential therapeutic target for increasing bacterial clearance during pneumonia-induced lung injury.

**Figure 7.**
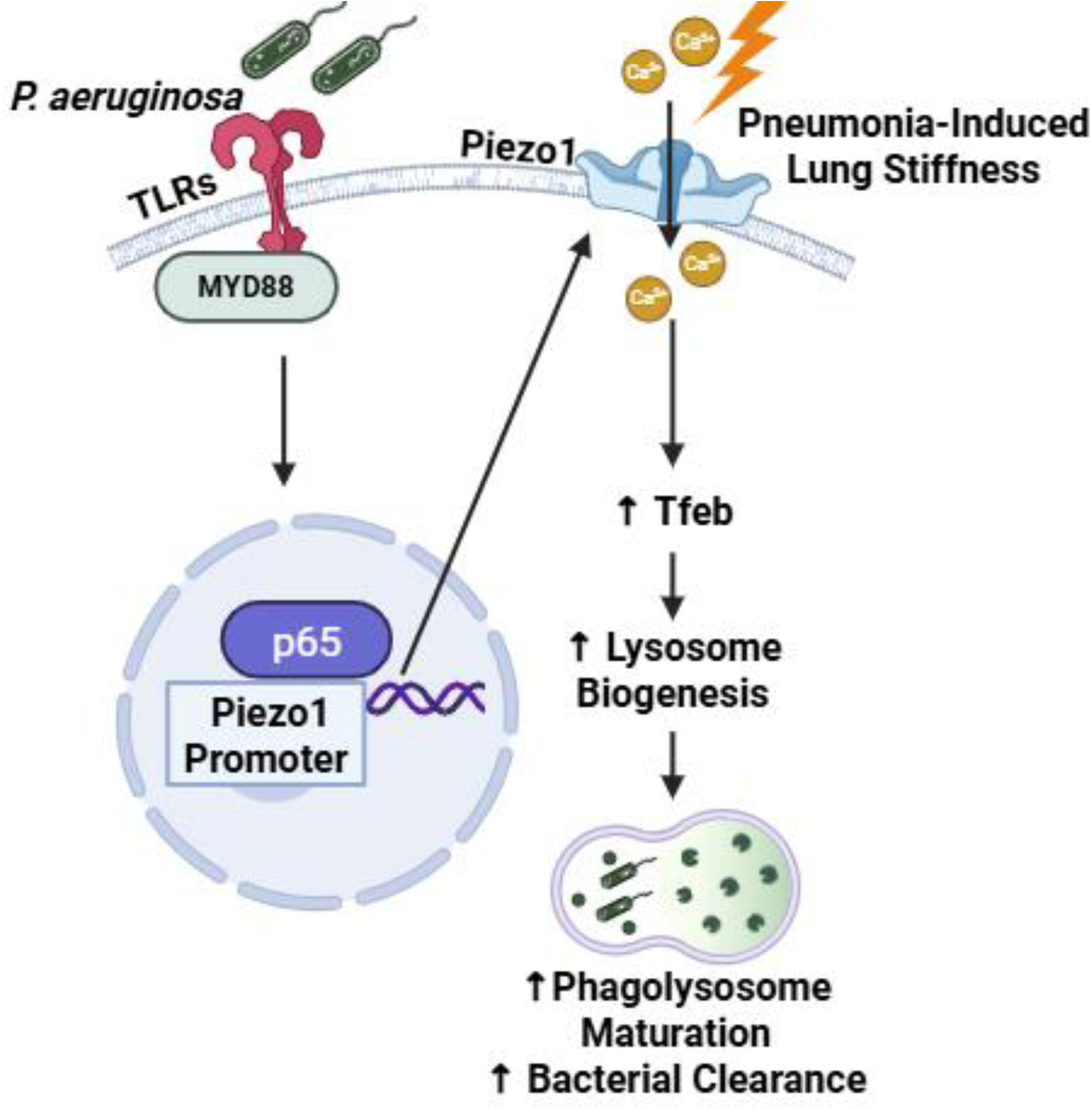
Schematic of the proposed mechanism of transcriptional regulation of Piezo1 by NF-κB/p65 during *P. aeruginosa* lung infection. Activation of TLRs by the respiratory pathogen, *P. aeruginosa* virulence factors (e.g., flagellin, LPS), activates NF-κB through MyD88 for increased Piezo1 transcription, surface abundance, and activation in response to injury-related increases in matrix stiffness. Piezo1 then augments Tfeb abundance which promotes lysosome biogenesis and phagolysosome function in macrophages, thereby augmenting bacterial clearance. Figure created with Biorender.com.

## Notes

Grant Support: This work was supported by R01HL155064 (RGS), R01HL158746 (MAO), R35GM149240 (VV), CFF ORSINI25Q0 (EMO), and SMARRT T32 which is funded by the National Institutes of Heart, Lung, and Blood grant, T32HL155005 (EMO).

### Competing Interest Statement

The authors have declared no competing interest.

