## Supplemental Figure 1 for "NF-κB-Dependent Transcriptional Regulation of Piezo1 Mediates Bacterial Clearance on Stiffened Lung Matrix"

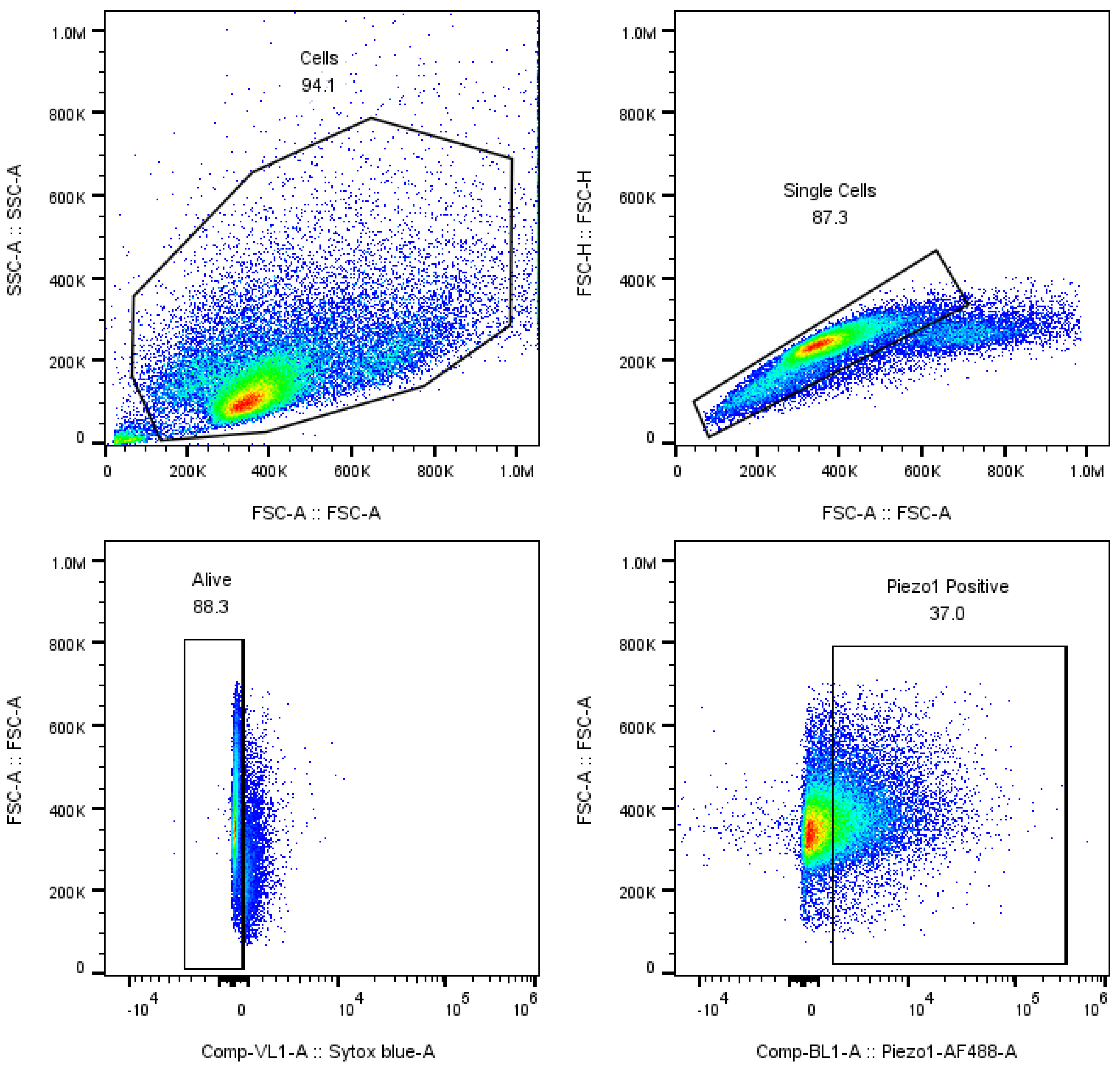
Cells
94.1
SSC-A :: SSC-A
FSC-A :: FSC-A
Single Cells
87.3
FSC-H :: FSC-H
FSC-A :: FSC-A
Alive
88.3
FSC-A :: FSC-A
Comp-VL1-A :: Sytox blue-A
Piezo1 Positive
37.0
FSC-A :: FSC-A
Comp-BL1-A :: Piezo1-AF488-A

**Supplemental Figure 1. Flow gating strategy** WT and MyD88$^{-/-}$ BMDMs were gated to remove debris, aggregates, and dead cells before gating on the frequency of Piezo1 positive cells.
