## Supplemental Figure 2 for "NF-κB-Dependent Transcriptional Regulation of Piezo1 Mediates Bacterial Clearance on Stiffened Lung Matrix"

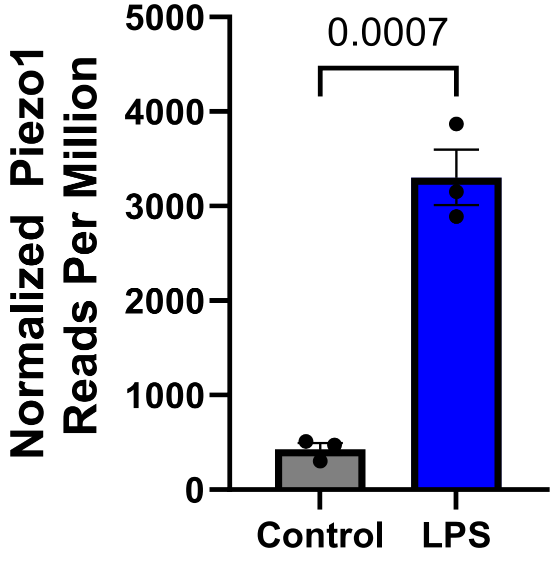
0.0007
Normalized Piezo1
Reads Per Million
5000
4000
3000
2000
1000
0
Control
LPS

**Supplemental Figure 2. LPS induces nascent Piezo1 mRNA production.** The first 4,000 base pairs of the Piezo1 gene in WT BMDMs ± LPS were quantified and graphed as normalized Piezo1 reads per million. Nascent mRNA strands are produced in BMDMs upon LPS stimulation according to published GRO-seq data (NCBI GEO No. GSE140611).
