## Supplemental Figure 3 for "NF-κB-Dependent Transcriptional Regulation of Piezo1 Mediates Bacterial Clearance on Stiffened Lung Matrix"

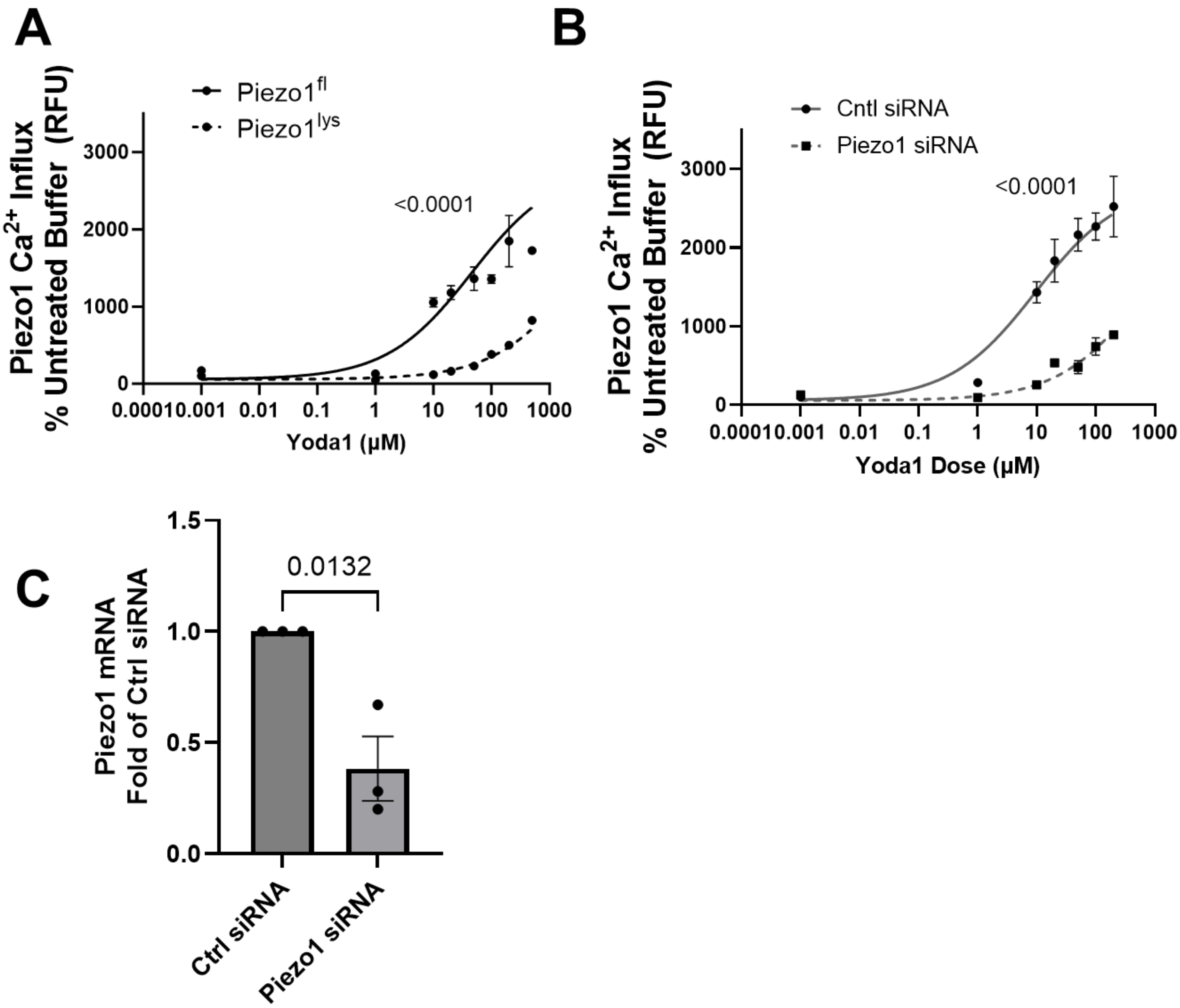
A
Piezo1^fl
Piezo1^lys
Piezo1 Ca2+ Influx
% Untreated Buffer (RFU)
<0.0001
3000
2000
1000
0
0.0001 0.001 0.01 0.1 1 10 100 1000
Yoda1 (μM)
B
Cntl siRNA
Piezo1 siRNA
Piezo1 Ca2+ Influx
% Untreated Buffer (RFU)
<0.0001
3000
2000
1000
0
0.0001 0.001 0.01 0.1 1 10 100 1000
Yoda1 Dose (μM)
C
Piezo1 mRNA
Fold of Ctrl siRNA
0.0132
1.5
1.0
0.5
0.0
Ctrl siRNA
Piezo1 siRNA

**Supplemental Figure 3. Piezo1 $Ca^{2+}$ channel activity is reduced with cre recombinase and Piezo1-targeted siRNA.** Piezo1 $Ca^{2+}$ channel activity was quantified in BMDMs with Piezo1 depletion by cre recombinase (Piezo1$^{LysMCre}$) or Piezo1-targeted siRNA. Piezo1 $Ca^{2+}$ channel activity in response to Yoda1 is reduced in (**A**) Piezo1$^{LysMCre}$ and (**B**) Piezo1 siRNA-treated BMDMs, demonstrating the specificity of Yoda1 for Piezo1. (**C**) Piezo1-targeted siRNA decreased Piezo1 mRNA by 2.6-fold, n = 3.
